# Role of anterograde motor Kif5b in clathrin-coated vesicle uncoating and clathrin-mediated endocytosis

**DOI:** 10.1101/284166

**Authors:** Yan-Xiang Ni, Wen-Qian Xue, Nan Zhou, Wing-Ho Yung, Rao-Zhou Lin, Richard Yi-Tsun Kao, Zhi-Gang Duan, Xu-Ming Tang, Meng-Fei Liu, Wen Zhang, Shuang Qi, Xi-Bin Lu, Jian-jiang Hu, Sookja Chung, You-Qiang Song, Jian-Dong Huang

## Abstract

Kif5b-driven anterograde transport and clathrin-mediated endocytosis (CME) are responsible for opposite intracellular trafficking, contributing to plasma membrane homeostasis. However, whether and how the two trafficking processes coordinate remain unclear. Here, we show that Kif5b directly interacts with clathrin heavy chain (CHC) at a region close to that for uncoating catalyst (Hsc70) and preferentially localizes on large clathrin-coated vesicles (CCVs). Uncoating *in vitro* is decreased for CCVs from the cortex of *kif5b* conditional knockout (mutant) mouse and facilitated by adding CHC-binding Kif5b fragments, while cell peripheral distribution of CHC or Hsc70 keeps unaffected by Kif5b depletion. Furthermore, cellular entry of vesicular stomatitis virus that internalized into large CCV is inhibited in cells by Kif5b depletion or introducing a dominant-negative Kif5b fragment. These findings showed a new role of Kif5b in CCV uncoating and CME, indicating Kif5b as a molecular knot connecting anterograde transport to CME.

## Introduction

Anterograde intracellular transport and endocytosis, two opposite trafficking processes, contribute to plasma membrane homeostasis that is fundamental to membrane integrity, cell survival, and function. Whether and how these two trafficking processes communicate remain unknown, although feedback mechanism was found to perceive and respond to changes in lipid abundance on the plasma membrane (Berchtold et al., 2012),.

Clathrin-mediated endocytosis (CME) is a conserved and efficient way of reducing protein levels on the plasma membrane and maintains normal cellular functions (McMahon and Boucrot, 2011, Ni et al., 2006, Conner and Schmid, 2003). It can also be hijacked by viruses, such as vesicular stomatitis virus (VSV), for entry into the host cells (Meertens et al., 2006, Cureton et al., 2009). The process of CME involves a cascade starting from the formation to the disassembly of clathrin-coated vesicles (CCVs). During CCV formation, clathrin triskelia are assembled into coat lattice surrounding the vesicle (Ferguson et al., 2007, Henne et al., 2010, Pechstein et al., 2010, Ehrlich et al., 2004). In contrast, the disassembly process is to release the triskelia from CCVs into cytosol, ensuring subsequent endocytic cellular events(Edeling et al., 2006). Heat shock cognate-70 protein (Hsc70) is an uncoating catalyst binding to clathrin on CCVs together with its cofactor Auxilin *in vitro* and *in vivo*, consequently leading to the collapse of clathrin lattice (Morgan et al., 2001, Xing et al., 2010, Yim et al., 2010). This disassembly/uncoating process is a critical step in CME and also regulated by other factors, such as synaptojanin(Harris et al., 2000) and endophilin(Gad et al., 2000, Milosevic et al., 2011). But more intrinsic regulators contributing to CCV uncoating under cellular physiological conditions remain to be clarified.

Anterograde transport driven by kinesin-1, which consists of two heavy chains and two light chains (KLCs), delivers various proteins to cell periphery along microtubules and increases their levels on the plasma membrane (Hirokawa et al., 2009). The conserved and ubiquitous kinesin-1 heavy chain Kif5b contains a microtubule-interacting motor, a stalk region, a KLC-binding site, and a cargo-binding tail (Hirokawa et al., 2009). Kif5b is essential for the transportation of membranous organelles and vesicles (Huang et al., 1999, Ong et al., 2000), including early endosomes (Loubery et al., 2008, Nath et al., 2007). After arriving at the cell periphery, Kif5b is released from microtubule and forms a folded and inactive conformation (Verhey and Hammond, 2009, Coy et al., 1999). However, it is unclear if the released Kif5b plays any unknown role around the plasma membrane, e.g. regulating endocytosis.

Here, we provide evidence that anterograde motor Kif5b binds to the proximal segment of clathrin heavy chain (CHC), localizes on relatively large CCVs and plays a noncanonical role in CCV uncoating without affecting the distribution of CHC or Hsc70 at the cell periphery. We evaluated the effects of Kif5b depletion on CME and found that Kif5b depletion interfered with large CCV mediated VSV cellular entry but hardly affects formation or function of synaptic vesicles. Furthermore, VSV entry was attenuated by applying a dominant-negative Kif5b fragment, which could overwhelm endogenous Kif5b for CHC binding *in vivo*. Overall, our study showed anterograde motor Kif5b plays a new role in CCV uncoating, hence regulating CME.

## Results

### Kif5b is associated with CHC and localizes on relatively large mouse cortical CCVs

To test whether Kif5b-mediated anterograde transport is linked to CME, we immuno-isolated Kif5b from mouse cortex and examined if any proteins involved in CME pathway were co-isolated. A ∼170 kDa band was repeatedly detected and subsequently identified by mass spectrometry as CHC (Figure 1A), which composes clathrin triskelion and is a major structural component of the CCV coat (Edeling et al., 2006, von Kleist et al., 2011).

**Figure 1.**
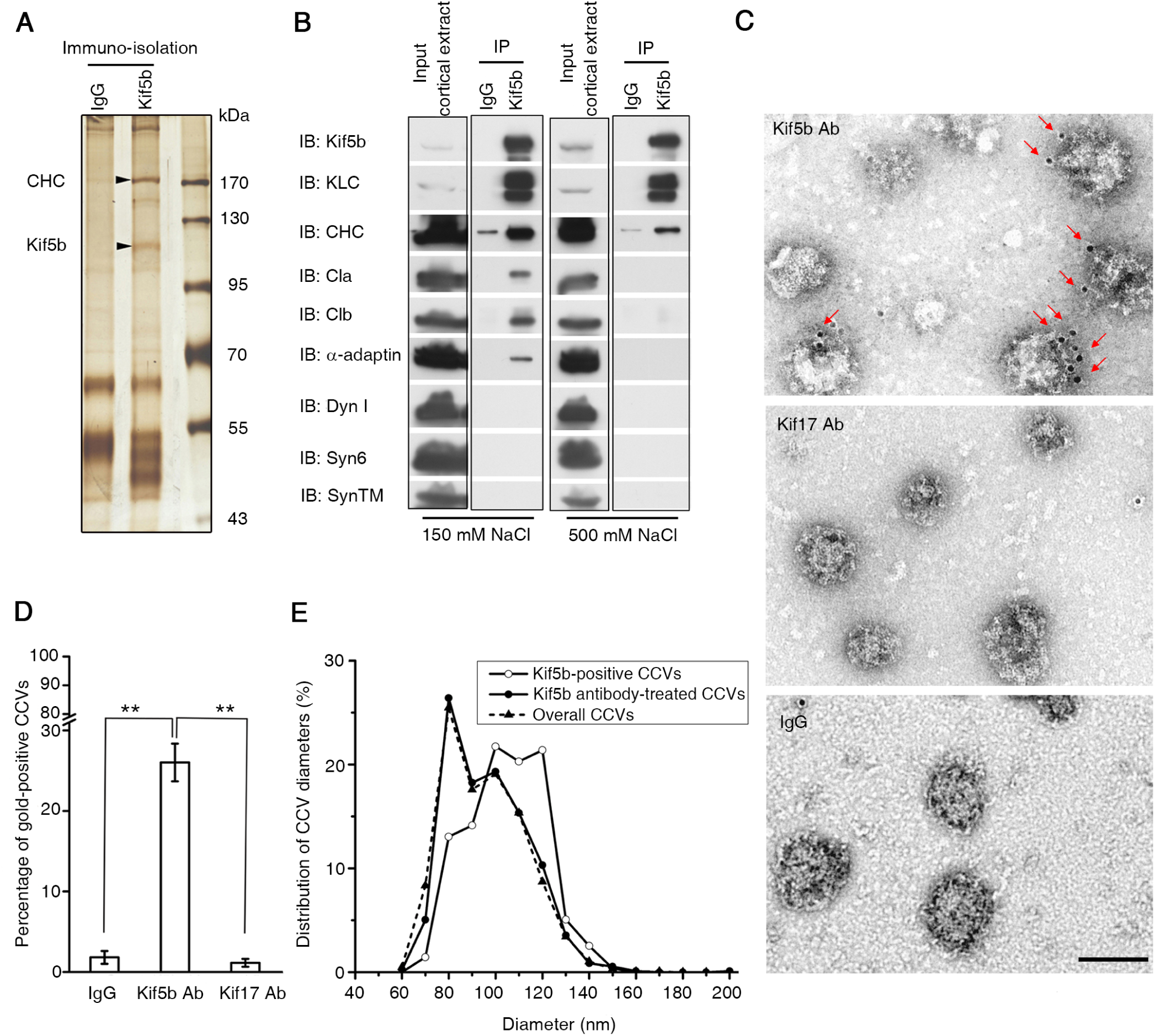
Kif5b is associated with CHC and localizes on relatively large mouse cortical CCVs. (**A**) Silver staining analysis of immune-isolated mouse cortical extracts by using Kif5b antibody or control IgG. The lower and upper band (arrowheads) were verified as Kif5b and CHC by mass-spectrometry. (**B**) Immunoprecipitation of Kif5b in mouse cortical extracts under 150 mM or 500 mM NaCl. The precipitates were analyzed by indicated antibodies. (**C**) Representative electron micrographs of cortical CCVs stained with normal IgG (bottom) or antibody against Kif5b (top) or Kif17 (middle), followed by incubation of 2^nd^ antibody conjugated to 10-nm gold particles. Arrowheads indicate the gold particles representing Kif5b on CCVs. Scale bar = 100 nm. (**D**) Percentage of gold-positive CCVs in different samples. 305, 500, and 324 CCVs from images of 35, 46, or 36 randomly-selected fields were analyzed for samples treated with Kif5b antibody, Kif17 antibody, or IgG, respectively. Error bars indicate s.e.m. \*\**P*<0.01. (**E**) Diameter distribution of analyzed CCVs for different groups. Data of 933 CCVs from Kif5b antibody-treated samples, 277 gold-positive CCVs, and 2733 overall CCVs were used for plotting. (IB: immunoblot; IP: immunoprecipitation; Cla: clathrin light chain A; Clb: clathrin light chain B; Dyn I: dynamin I; Syn6: syntaxin 6; SynTM: synaptotagmin)

To confirm the association of CHC with Kif5b, we immunoprecipitated Kif5b from mouse cortex under both physiological (150 mM) and a more stringent (500 mM) salt condition, and probed the co-precipitates by different antibodies. In addition to detecting KLC, we observed CHC in precipitates (Figure 1B). Kif5b failed to co-precipitate syntaxin-6, synaptotagmin, or dynamin I, which were present in the cortical input, indicating the specificity of the co-immunoprecipitation assay. Notably, besides CHC, other CHC-associated coat components, such as α-adaptin, was also detected (Figure 1B). α-adaptin is a specific subunit of AP2 adaptor complex that binds to CHC and localizes on endocytic CCVs rather than non-endocytic CCVs traveling between *trans*-Golgi and lysosome(Kelly et al., 2014). Thus the co-precipitation of CHC and α-adaptin indicated the association of Kif5b with endocytic CCVs.

To directly address if Kif5b localizes on cortical CCVs, we purified CCVs from mouse cortex and confirmed them by electron microscopy (EM) as electron-dense particles with characteristic CCV structures and diameter distribution (Howe et al., 2001, Fischer et al., 1999, Maycox et al., 1992) (Figure S1). Then we performed immunogold EM with the purified CCVs using different primary antibodies and a gold-conjugated secondary antibody. Gold particles were observed specifically associated with CCVs treated with Kif5b antibody, rather than those with Kif17 antibody or control IgG (Figure 1C and 1D). The localization of Kif5b on endocytic CCVs is likely relying on the specific interaction between Kif5b and CHC or CHC-binding proteins, as gel-enhanced liquid chromatography/tandem MS analysis (GeLC-MS/MS) of the cortical precipitates by Kif5b antibody detected CHC and CHC-associated proteins rather than other CME cargo proteins (e.g., transferrin receptor) or proteins on non-endocytic CCVs (e.g., AP1, AP3, and AP4) (Table S1). Interestingly, when measuring the diameters of CCVs, we found that Kif5b was mainly localized on the CCVs of relatively large size (a diameter peak > 90 nm), while overall CCVs analyzed or the population of CCVs treated with Kif5b antibody showed a diameter peak at around 80 nm (Figure 1E). Taken together, these results demonstrate specific localization of anterograde motor Kif5b on cortical CCVs, particularly the large ones.

### Kif5b directly interacts with CHC proximal segment via a 25-amino acid tail region

To map the potential binding site on Kif5b for clathrin, we generated a series of Kif5b fragments that were fused to glutathione S-transferase (GST) (Figure 2A). The fusion proteins were then immobilized and incubated with mouse cortical extracts. CHC and α-adaptin were specifically pulled down by the Kif5b C-terminal fragment (residues 679-963) rather than the motor (residues 1-413) or stalk region (residues 414-678) (Figure 2B). The C-terminal fragment was further divided into two parts, Kif5b^679-849^ and Kif5b^850-963^. CHC and α-adaptin were pulled down by Kif5b^850-963^, but not Kif5b^679-849^ containing a KLC-binding site (Figure 2B), indicating a Kif5b-CHC association independent of KLC or KLC-Hsc70 interaction(Terada et al.). Further experiments with shorter truncations revealed that the tail (Kif5b^891-963^), Kif5b^850-915^ and Kif5b^891-935^ fragments were able to pull down CHC or α-adaptin, whereas Kif5b^850-890^ fragment could not (Figure 2C), indicating that the CHC-binding site was localized within the 891-915 region of Kif5b. Indeed, it was found that the fragment Kif5b^891-915^ was sufficient to pull down CHC and α-adaptin (Figure 2D). Collectively, these data revealed that the 25-amino acid tail region (residues 891-915) of Kif5b was a CHC-binding site and that Kif5b-CHC interaction was independent of KLC. We also studied the interaction between Kif5b^891-963^ and different recombinant fragments of CHC (Figure 2E) and observed a direct interaction between CHC^1280-1429^, the N-terminus of CHC proximal segment (Fotin et al., 2004b, Xing et al., 2010), and Kif5b tail (Figure 2F and 2G). Taken together, these data demonstrated a direct interaction between residues 891-915 of Kif5b and N-terminus of the proximal segment of CHC.

**Figure 2.**
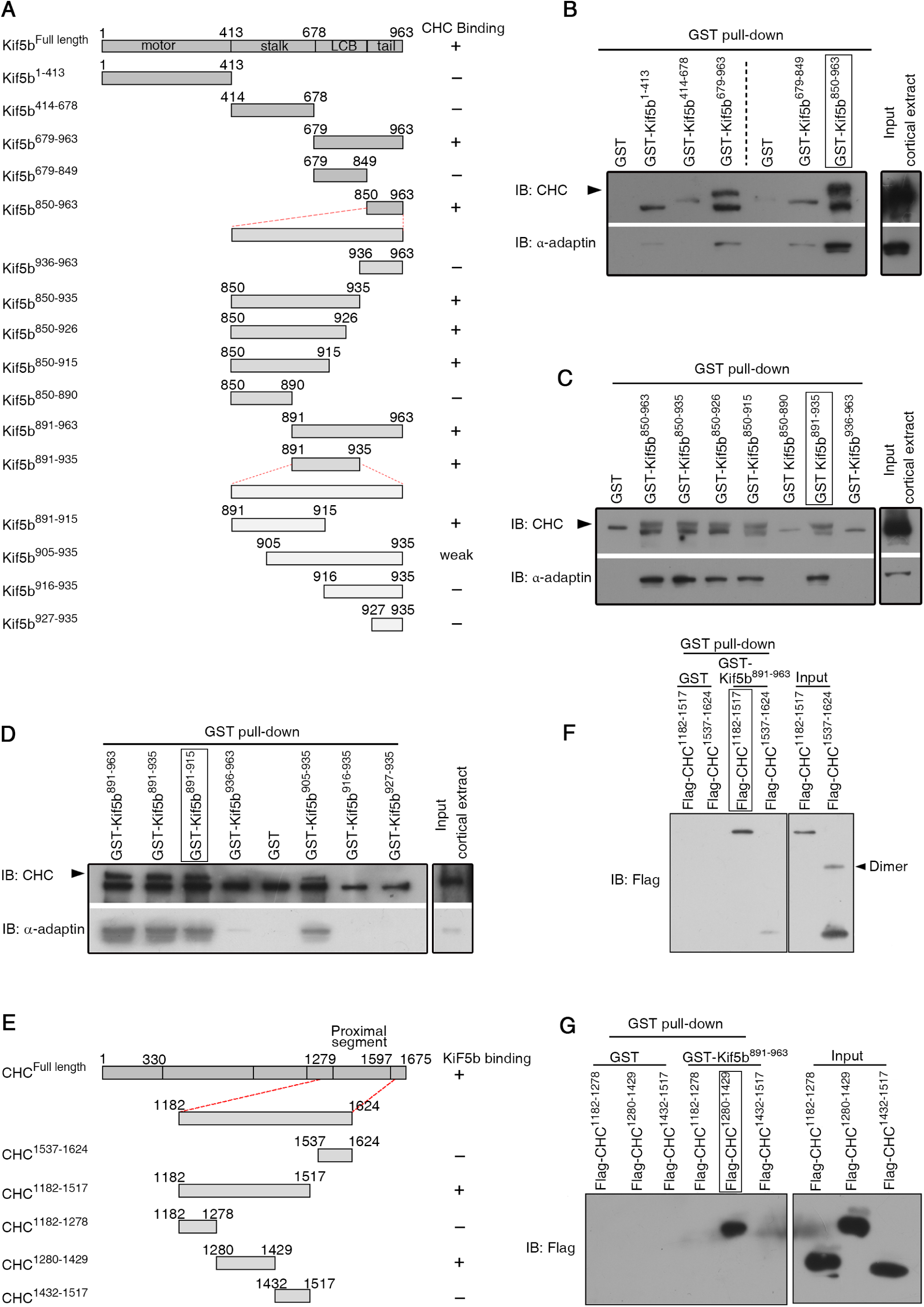
Kif5b directly interacts with CHC proximal segment via a 25-a.a tail region. (**A**) Schematic diagram of Kif5b fragments and their ability to bind CHC. LCB stands for light chain binding region. (**B-D**) Pull down of CHC and α-adaptin in mouse cortices by different GST-fused Kif5b deletions (**B**) or tail truncations (**C** and **D**). Arrowheads indicates CHC above the non-specific band. (**E**) Schematic diagram of CHC fragments and their ability to bind Kif5b tail. (**F-G**) Pull down of recombinant flag-tagged CHC fragments from bacterial lysates by Kif5b^891-963^.

### Kif5b plays a role in regulating uncoating of mouse cortical CCVs without affecting the cell peripheral distribution of CCV coat proteins and uncoating catalyst

Given that Kif5b interacts with CHC and localizes on CCVs, we wondered if this well-known anterograde motor is responsible for the intracellular trafficking of components of endocytic CCVs. To address this question, we knocked down Kif5b in Neuro2a cells with the use of shRNA (Figure S2A and S2B). Immunofluorescence staining and quantitative analysis revealed that the abundance of CHC, α-adaptin and also Hsc70 in cellular compartments close to plasma membrane remained almost unchanged in Kif5b-depleted cells (Figure 3A and 3B). These results suggest that cell peripheral distribution of major CCV components and uncoating catalyst is likely independent of Kif5b.

**Figure 3.**
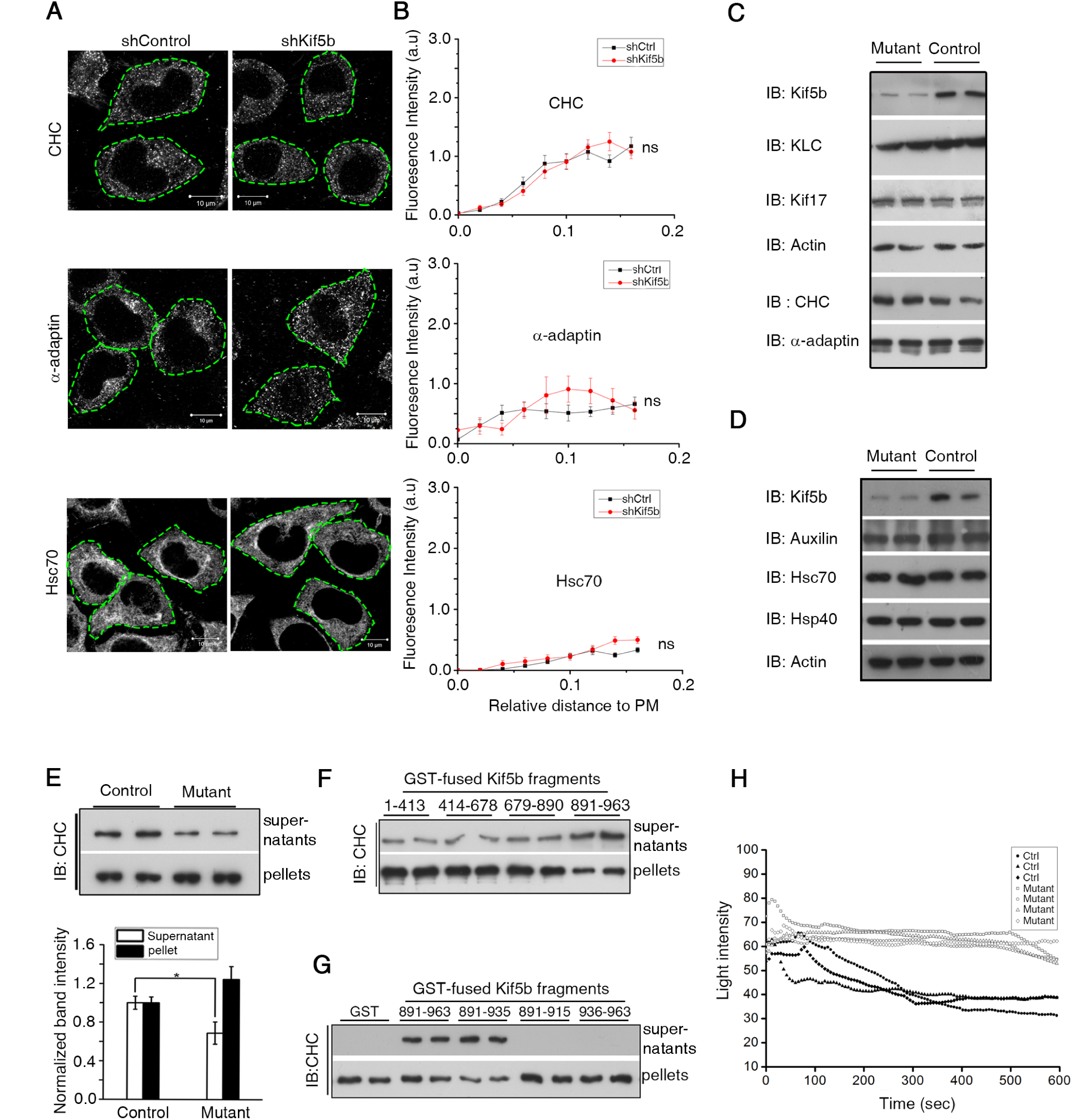
Kif5b regulates CCV coating without affecting cell peripheral distribution of CCV coat proteins and uncoating catalyst. (**A**) Representative immunofluorescence staining images for CHC, α-adaptin and Hsc70 in shKif5b or shControl RNA treated Neuro2a cells. Green dotted lines delineate the cell surface defined by co-staining of F-actin with phalloidion. Scale bar =10 µm. (**B**) Quantitative analysis of subcellular distribution of CHC, α-adaptin and Hsc70. The distribution was quantified by measuring the line profile of fluorescence intensity from the plasma membrane (PM) to the nucleus (See also Figure S5). Error bars indicate s.e.m. n=30. ns, not significant (*P* > 0.05). (**C** and **D**) Representative Western blots of cortical proteins or uncoating related proteins in cortices of *kif5b* mutant mice or control littermates. (**E**) Up: Uncoating assay of CCVs from mutant or control mouse cortices. The amount of CHC released into supernatants was representative of uncoating. Bottom: Quantitative analysis of released CHC. Error bars indicate s.e.m. n=3. \**P*<0.05. (**F** and **G**) Uncoating assay of CCVs treated with or without different recombinant fragments of Kif5b. (**H**) Representative disassembly curve of purified cortical CCVs from mutant or control mice detected by light scattering assay *in vitro*.

To investigate the potential new function(s) of Kif5b on CCVs, we employed a *kif5b* conditional knockout (mutant) mouse (*Camk2a-cre; kif5b*^*-/floxP*^, Figure S2C) that expressed Cre in the forebrain (Tang et al., 1999). In cortices of mutant mice, Kif5b expression was decreased by more than 60% compared with controls, whereas other kinesin proteins (e.g., KLC and Kif17) remained almost unchanged (Figure 3C and S3A). We also examined the expressions of CCVs proteins and found that levels of coat proteins (e.g., CHC and α-adaptin) and uncoating-related proteins (Hsc70, cofactors Hsp40, and Auxilin) were not affected in cortices of mutant mice (Figure 3D and S3B). These findings implied that Kif5b is likely involved in other aspects rather than anterograde transport of CCV proteins.

We then asked whether endogenous Kif5b is involved in CCV uncoating under physiological conditions. To this aim, we modified an established centrifugation-based uncoating assay (Hannan et al., 1998, Barouch et al., 1994). After ultracentrifugation, disassembled CCVs released coat proteins into the supernatant, whereas intact CCVs and uncoated vesicles remained in the pellet. Significantly less CHC was released into the supernatant from mutant CCV reactions than that from control reactions, even though the input CCVs were of the same amount (Figure 3E), indicating Kif5b was involved in CCV uncoating. Next, we added different GST-fused Kif5b fragments (Figure S4A) into uncoating reaction. Addition of Kif5b tail that carries the CHC-binding site was able to increase CHC release, but not for other fragments without CHC-binding site, including the motor (Kif5b^1-413^), stalk region (Kif5b^414-678^), and KLC-binding site-containing fragment (Kif5b^679-890^) (Figure 3F). In addition, the Kif5b tail fragment Kif5b^891-963^ could cause a dose-dependent increase of CHC release in the uncoating reaction (Figure S4B and S4C). These results indicated the requirement of Kif5b-CHC interaction for uncoating. We further analyzed the effects of various tail regions and observed that uncoating was facilitated by Kif5b^891-935^ similar to the tail Kif5b^891-963^ whereas the minimal CHC-binding region Kif5b^891-915^ was not able to facilitate uncoating (Figure 3G and S4D). Notably, the requirement of Kif5b-CHC interaction for uncoating was further proved by using a real-time uncoating assay of light scattering intensity that dynamically reflects vesicle diameter change (Rothnie et al., 2011, Young et al., 2013). We found that CCVs from mutant mouse cortices exhibited a slower decrease of light scattering intensity than those from control samples during the 10 min reaction period (Figure 3H). Taken together, these data showed that anterograde motor Kif5b plays a role in regulating the uncoating of mouse cortical CCVs.

### Large CCV-mediated endocytosis of VSV is inhibited by Kif5b depletion or a dominant-negative Kif5b fragment for uncoating

Since Kif5b participates in CCV uncoating, a critical step in CME, we next address if Kif5b is involved in regulating CME. Although many cell-surface receptors and virus enter cells via CME, we adopted VSV cellular entry as our model here, because Kif5b mainly localizes on large CCVs and VSV internalization involves large CCVs with a dimension of about 180×70 nm during infection (Cureton et al., 2010). Then we applied replication-deficient VSV pseudoparticles and infected cells for different time slots, followed by immunofluorescence staining of all internalized virus in cells with the use of a VSV-specific antibody (Figure 4A). Quantitative analysis showed a remarkable accumulation of internalized VSVs along infection time in shControl cells (Figure 4B, black bars) but not in shKif5b cells (Figure 4B, white bars). A significant difference was observed between the two groups after 15 min infection (Figure 4B). In contrast, when examining synaptic vesicles, the endocytosis of which involves CCVs of small size (40-45 nm in diameter) (Zhang et al., 1998, Watanabe et al., 2014), no changes of synaptic vesicle formation or related function were observed by *kif5b* knockout in mouse cortex (Figure S6). Taken together, these data showed that Kif5b regulates large CCV-involved cellular entry of VSV, consistent with the localization and uncoating function of Kif5b on CCVs.

**Figure 4.**
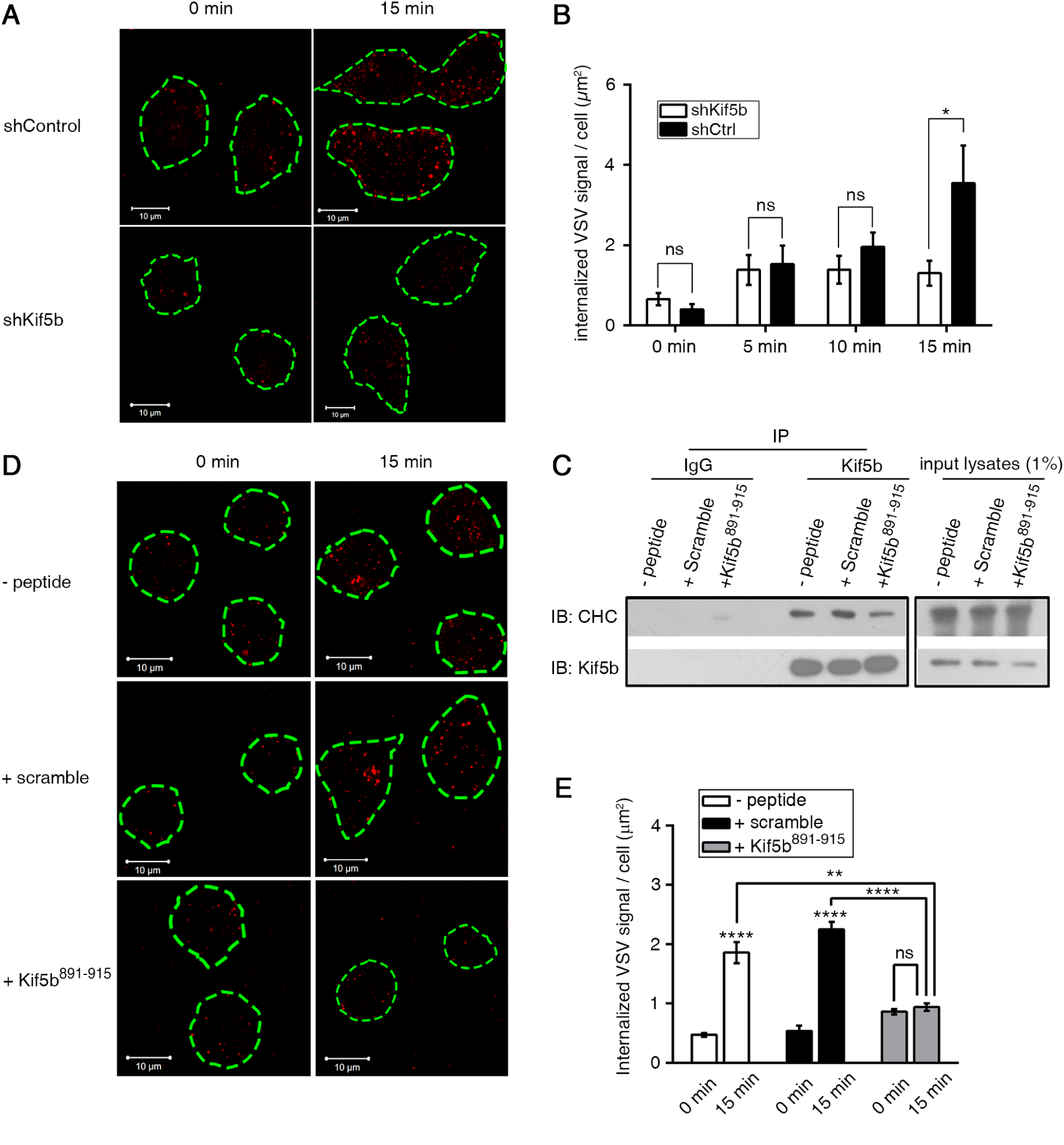
Kif5b regulates large CCV-dependent VSV cellular entry. (**A**) Representative immunofluorescence staining images of internalized VSVs in shKif5b or shControl Neuro2a cells after 0 or 15 min VSV infection. (**B**) Quantitative analysis of internalized VSV signals for shKif5b or shControl Neuro2a cells infected by VSV with different time slots. (**C**) Co-immunoprecipitation of CHC with Kif5b from Neuro2a cells with 1 h pre-treatment with or without 100 μM cell-penetrating scramble or Kif5b^891-915^ peptide at 37 °C. (**D**) Representative immunofluorescence staining images of internalized VSVs in Neuro2a cells with 15 min infection after 1 h pretreatment without peptide or with 100 μM scramble or Kif5b^891-915^ peptide at 37 °C. Green dashed lines delineate the cell periphery defined by co-staining of F-actin with phalloidion. Scale bar =10 μm. (**E**) Quantitative analysis of internalized VSV signal in different Neuro2a cell groups indicated in **(D)**. Error bars indicate s.e.m. n>30. \*\*\*\**P*<0.0001; ** *P*<0.01; \**P*<0.05. ns, not significant (*P* > 0.05).

To further confirm the aforementioned regulation of VSV entry is contributed by new role of Kif5b in CCV uncoating, we applied a TAT-conjugated (Jones et al., 2005) cell-penetrating peptide corresponding to Kif5b^891-915^, which was shown to bind to CHC without facilitating CCV uncoating *in vitro* (Figure 2D and 3G) and is expected to act as a dominant-negative fragment for uncoating *in vivo*. A dominant-negative approach typically leads to loss of function of a wild-type protein through the sequestration of its effectors. Consistently, we found that less CHC was co-immunoprecipitated with Kif5b in cells pretreated with Kif5b^891-915^ peptide than in controls without any peptide pre-treatment or with a scramble peptide with the same amino acids composition as Kif5b^891-915^ (Figure 4C), suggesting that Kif5b^891-915^ could overwhelm endogenous Kif5b for CHC binding *in vivo*. We then assessed the effect of peptide pretreatment on VSV cellular entry. Immunofluorescence staining and quantitative analysis of internalized VSVs in cells with 15 min VSV infection showed a significant decrease of VSV entry in the cell group pretreated with Kif5b^891-915^ peptide compared with the control groups (Figure 4D and 4E). Overall, these data show that Kif5b regulates VSV cellular entry depending on its role in uncoating.

## Discussion

Various mechanisms contribute to the maintenance of plasma membrane homeostasis, including coordination between different endocytic pathways (Pelkmans et al., 2005). In this study, we observed the localization of Kif5b on large CCVs at the N-terminal part of CHC proximal segment, which is spatially near the Hsc70-binding tripod on CCVs (Fotin et al., 2004b, Fotin et al., 2004a, Xing et al., 2010). Depletion of Kif5b decreased CHC-Hsc70 interaction and impeded CCV uncoating that is essential for CME. Therefore, Kif5b may serve as a molecular knot, contributing to the connection between anterograde transport and CME, and also the potential feedback loop in cells for plasma membrane homeostasis. These findings provide a possible explanation for how cells can partially protect plasma membrane integrity against anterograde transport fluctuations: as kinesin-1-mediated anterograde transport delivers more proteins to the plasma membrane, more Kif5b will reach the cell periphery to increase CME; conversely, when kinesin-1-mediated anterograde transport slows down, less Kif5b is available to CCVs, thus decreasing CME.

CCV uncoating have been intensively studied by using recombinant clathrin cages generated *in vitro*, especially when dissecting the mechanism of how a known uncoating regulator, such as Hsc70, conveys its effect (Rothnie et al., 2011, Fotin et al., 2004b, Xing et al., 2010). Here, to address Kif5b as a new intrinsic uncoating regulator, we applied purified cortical CCVs instead of recombinant clathrin cages and chose not to add any extra factors like Hsc70 or Auxilin into the reaction. This approach maintained the ratio of different proteins on the assayed CCVs largely similar to that under physiological conditions, which avoided masking the undetermined function of Kif5b. Given the close binding sites on CHC for Kif5b and Hsc70, the unchanged cell peripheral distribution and unaltered expression levels of CHC and Hsc70 in Kif5b mutant samples, we speculate that Kif5b contributes to uncoating via stabilizing or reinforcing the interaction between CHC and Hsc70 on CCVs rather than recruiting Hsc70 to CCVs. Taken together, this study enriches our knowledge of uncoating regulation under cellular physiological conditions.

Kif5b was found to preferentially localize on relative large CCVs with an external diameter more than 90 nm (Figure 1E). Such preference is possibly due to the different formation process for large and small CCVs(Cureton et al., 2010, Boucrot et al., 2006).In line with this observation, Kif5 depletion could specifically interfere large CCV-dependent VSV uptake (Figure 4A and 4B) rather than the formation and function of synaptic vesicle (Figure S6), the internalization of which relies on CCVs of relatively small size. In our study, introducing a 25-amino acid dominant-negative Kif5b fragment in uncoating decreased VSV cellular entry (Figure 4D and 4E), indicating the potential application of CHC-binding peptide in impeding VSV infection. Therefore, the Kif5b-CHC interaction might serve as a target for developing anti-viral drugs without side effects in normal cellular functions such as synaptic transmission.

## Methods

### CCV purification

CCVs were purified from mouse cortices according to modified protocols (Howe et al., 2001, Fischer et al., 1999, Maycox et al., 1992). Mice cortices were homogenized in Mes buffer (100 mM MES, 1 mM EGTA, 2 mM MgCl_2_, pH 6.5) and centrifuged at 1,000 × g for 10 min at 4 °C. The resultant supernatant was layered onto a 5% glycerol pad and centrifuged at 100,000 × g for 1 h. The pellet was re-suspended, and mixed 1:1 with Mes buffer containing 12.5% Ficoll 400 and 12.5% sucrose and centrifuged at 40,000 × g for 40 min. Supernatant was diluted 1:5 with Mes buffer, filtered in 0.2-μm filters (Corning, 431224), and centrifuged at 33,000 × g for 1 h. The CCV-containing pellet was re-suspended for further use.

### Centrifugation-based CCV uncoating

The uncoating assay was modified from established protocols (Hannan et al., 1998, Barouch et al., 1994). The CCVs (120 μg) were incubated for 8 min at 25°C in the absence or presence of the indicated fusion proteins (20 μg) in a total volume of 500 μl containing 2 mM ATP, 75 mM KCl, 5 mM MgCl_2_, and PIC. The reaction mixtures were centrifuged at 100,000 × g for 10 min at 4 °C. Supernatants and pellets were subjected to Western blot analysis.

### Light scattering-based CCV uncoating

Equal amount of purified CCVs from mutant or control mice was mixed at 4 °C with 1 mM ATP in a total volume of 30 μL in buffer (40 mM HEPEs, pH 7.0, 75 mM KCl, 4.5 mM Mg acetate). The mixtures were added to 384-well black polystyrene plate (Sigma, CLS3702-25EA) on the ice and spun down for 5 sec to remove any bubbles inside the reaction. Light scattering was immediately monitored using Wyatt DynaPro at 25 °C every 3 s for up to 600 s (Rothnie et al., 2011, Young et al., 2013).

### VSV pseudoparticle infection and analysis

Aliquots of viral dilutions produced from transfection of HEK293T cells with VSV pseudoparticle packaging vectors were incubated with Neuro2a cells for 1 h at room temperature (22 °C). After washing by DMEM, cells were shifted to 37 °C for indicated time for virus infection. Internalized VSVs were fixed and detected by an antibody against VSV-G (Abcam, ab1874). Intracellular VSV signals were blindly analyzed by the Particle Analyze function of ImageJ with a uniformly defined image threshold. Background signal noise calculated from cell groups without VSV treatment was subtracted.

## Acknowledgments

We would like to thank S. Brady and G. Morfini for the pCDNA3.Kif5b plasmid, D. Jin and J. Ng for the modified pLL3.7/U6 promoter vector, J. Garcia and M. Kudelko for the packaging plasmids for VSV pseudoparticles, and G.Niu for supporting in data analysis. We also thank Y. Guan, H. Zhu, and S. Xiu for their helpful advice and support. We thank electron microscopy unit, core facility center, and center of genomic sciences at the University of Hong Kong for their technical support. This work was supported by grants from the Hong Kong Research Grants Council (HKU 768113M) and the University of Hong Kong Small Project Fund to J.D.H. and Y.X.N, and partially by a National Basic Research Program of China (973 Program, 2014CB745200) from the Ministry of Science and Technology of PRC, and by Shenzhen Peacock project (KQTD2015033117210153); the Shenzhen Science and Technology Innovation Committee Basic Science Research Grant (JCYJ20150629151046896).

## Author contributions

J. Huang and Y. Ni designed the research. Y. Ni, WQ Xue and N. Zhou performed the experiments and wrote the manuscript. W. Yung contributed to the synaptic transmission experiments. R. Lin provided constructs of Kif5b truncations. R. Kao contributed to the light scattering-based uncoating assay. M. Liu and X. Tang provided the shRNA lentivirus for Kif5b knockdown. W. Zhang provided constructs of CHC fragments. Q. Shuang contributed to purification of CCVs. Z. Duan contributed to mouse breeding.

