## Supplementary Materials for "Role of anterograde motor Kif5b in clathrin-coated vesicle uncoating and clathrin-mediated endocytosis"

### Figure S1

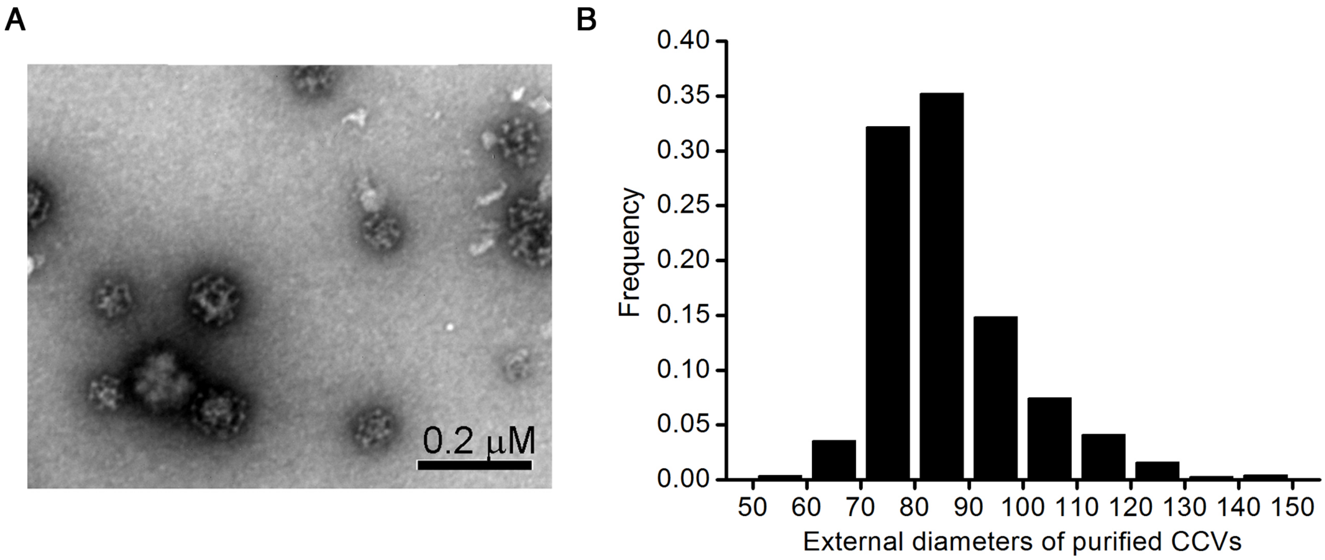
A
0.2 μM
B
Frequency
0.00
0.05
0.10
0.15
0.20
0.25
0.30
0.35
0.40
50 60 70 80 90 100 110 120 130 140 150
External diameters of purified CCVs

**Figure S1. Confirmation of the purified cortical CCVs by negative staining and electron microscopy, related to Figure 1**

(**A**) CCVs were purified from mice cortices and examined by electron microscopy after negative staining. The purified CCVs preserve their typical coat structures well with application of the ice-cold acidic buffer. (**B**) External diameter distribution of purified cortical CCVs measured from high-magnification electron micrographs. Scale bar = 0.2 μm.

### Figure S2

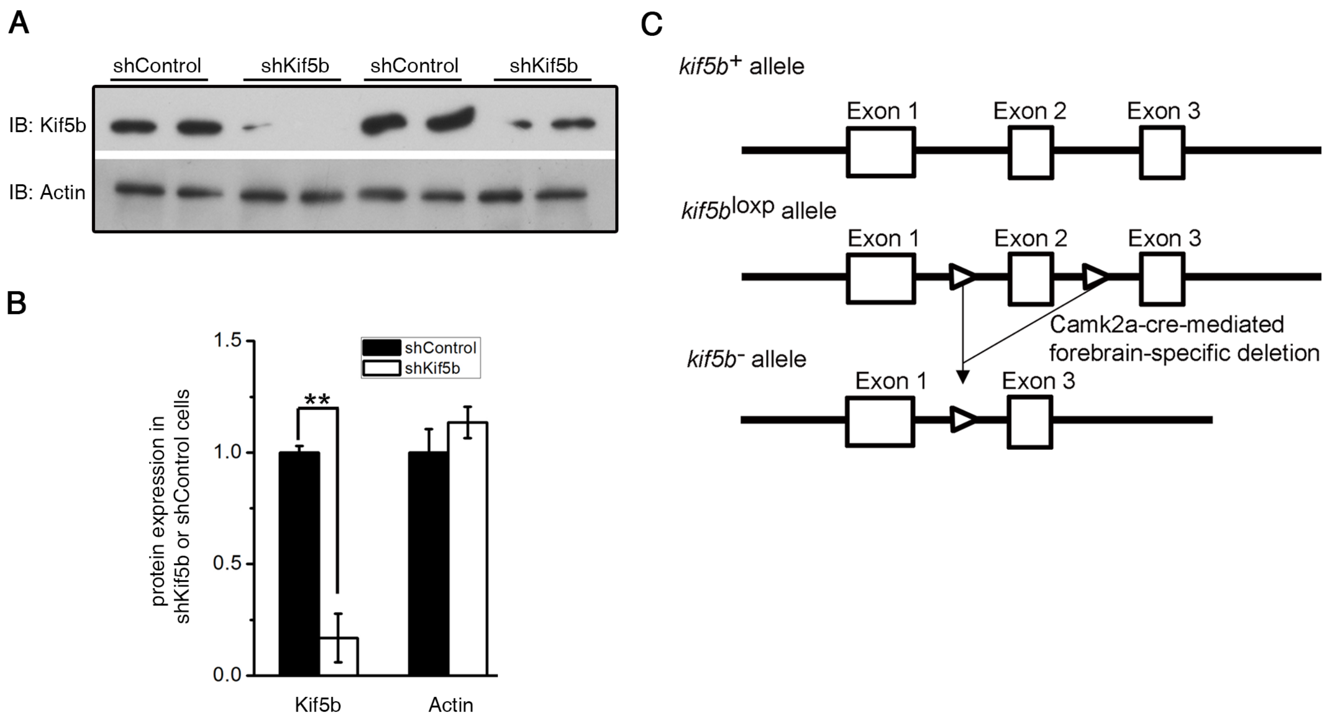

A
shControl
shKif5b
shControl
shKif5b
IB: Kif5b
IB: Actin
B
protein expression in shKif5b or shControl cells
1.5
1.0
0.5
0.0
shControl
shKif5b
**
Kif5b
Actin
C
kif5b+ allele
Exon 1
Exon 2
Exon 3
kif5bloxp allele
Exon 1
Exon 2
Exon 3
Camk2a-cre-mediated forebrain-specific deletion
kif5b- allele
Exon 1
Exon 3

**Figure S2.** ***kif5b*** **knock-down by shRNA and** ***kif5b*** **conditional knock-out in mouse cortex, related to Figure 2.**

(**A**) Kif5b depletion in Neuro2a cells by shRNA was confirmed by Western blot analysis with indicated antibodies.
(**B**) Quantitative analysis of the indicated bands in left (**A**). The protein expression levels in shControl cells were normalized to 1.0, and the levels of the proteins in shKif5b samples were expressed as the relative to the normalized values. Error bars indicate s.e.m. **P<0.01 as analyzed by Student's t-test.
(**C**) Schematic illustration of Camk2a-cre-mediated deletion of exon 2 from the *kif5b* allele.

### Figure S3

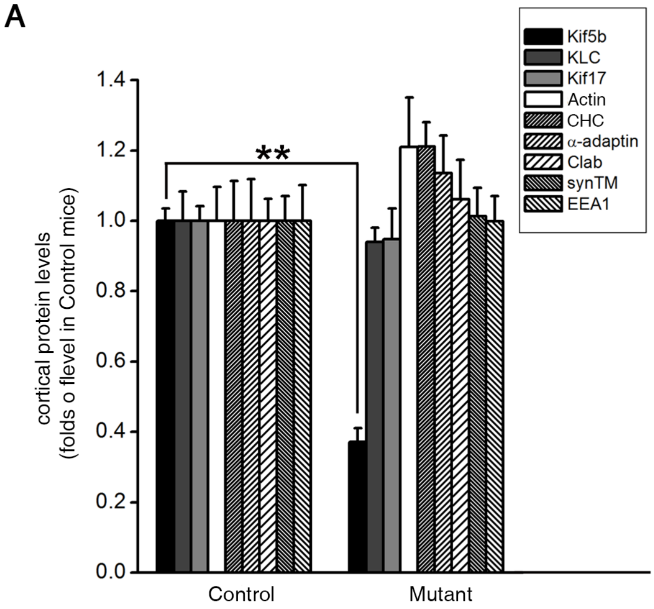
A
Kif5b
KLC
Kif17
Actin
CHC
α-adaptin
Clab
synTM
EEA1
1.4
1.2
1.0
0.8
0.6
0.4
0.2
0.0
**
cortical protein levels
(folds o flevel in Control mice)
Control
Mutant

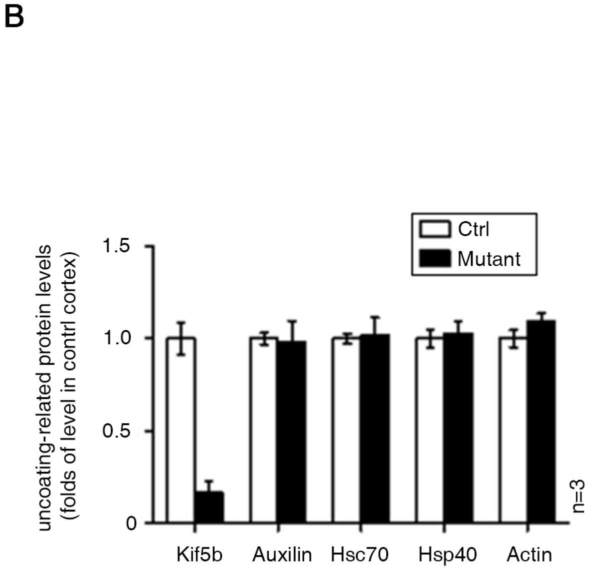
B
Ctrl
Mutant
1.5
1.0
0.5
0
uncoating-related protein levels
(folds of level in contrl cortex)
n=3
Kif5b
Auxilin
Hsc70
Hsp40
Actin

**Figure S3. Quantitative analysis of cortical protein and uncoating related protein levels in the mutant or control mouse cortex, related to Figure 3.**

(**A**) Quantitative analysis of the indicated cortical protein levels (Figure 3C) from kif5b mutant (n= 4) and control (n= 4) littermates. The levels of the indicated proteins in the controls were normalized to 1.0 and the levels of the proteins in mutant samples were expressed as the relative to the normalized values. Error bars indicate s.e.m. **P<0.01 as analyzed by Student's t-test.
(**B**) Quantitative analysis of the indicated uncoating related protein levels (Figure 3D) from kif5b mutant (n= 3) and control (n= 3) littermates. The levels of the indicated proteins in the controls were normalized to 1.0 and the levels of the proteins in mutant samples were expressed as the relative to the normalized values. Error bars indicate s.e.m. **P<0.01 as analyzed by Student's t-test.

### Figure S4

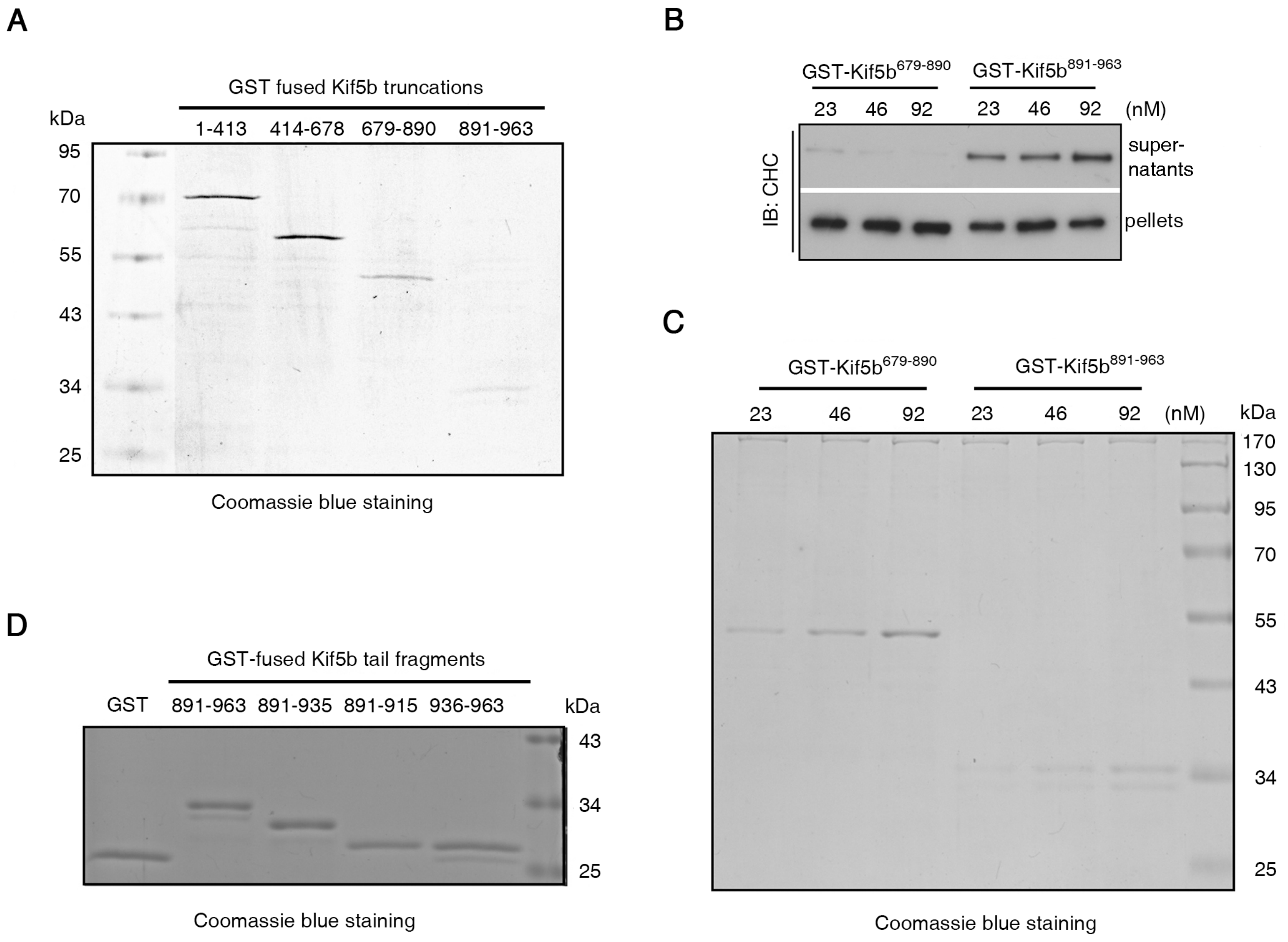

A
GST fused Kif5b truncations
kDa
1-413
414-678
679-890
891-963
95
70
55
43
34
25
Coomassie blue staining
B
GST-Kif5b$^{679-890}$
GST-Kif5b$^{891-963}$
23 46 92 23 46 92 (nM)
IB: CHC
super-
natants
pellets
C
GST-Kif5b$^{679-890}$
GST-Kif5b$^{891-963}$
23 46 92 23 46 92 (nM)
kDa
170
130
95
70
55
43
34
25
Coomassie blue staining
D
GST-fused Kif5b tail fragments
GST
891-963
891-935
891-915
936-963
kDa
43
34
25
Coomassie blue staining

**Figure S4. Kif5b tail fragment 891-963 caused a dose-dependent increase of CHC uncoating and GST-fused Kif5b fragments for CCV uncoating assay, related to Figure 3.**

(**A**) Coomassie blue staining of GST-fused Kif5b fragments used in the uncoating assay in Figure 3F.
(**B**) Increasing concentrations of Kif5b tail (891-963) or the fragment containing the KLC-binding site (residues 670-890) were applied in vitro in the uncoating assay. CHC released into supernatants or remaining in the pellets was determined by Western blot analysis.
(**C**) Coomassie blue staining of GST-fused Kif5b fragments used in the uncoating assay in Figure S4B.
(**D**) Coomassie blue staining of GST-fused Kif5b fragments used in the uncoating assay in Figure 3G.

### Figure S5

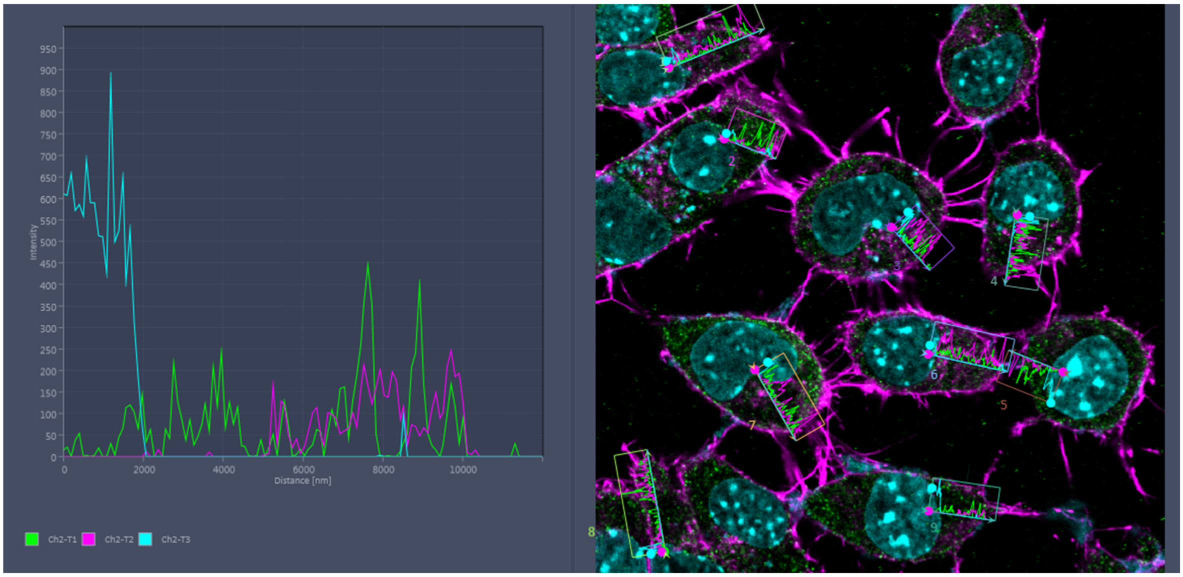

**Figure S5. Representative measurements of subcellular distribution of CHC in neuro2a cells, related to Figure 3.**

Images were acquired by a Zeiss LSM780 confocal laser scanning microscope. The fluorescence profile was measured by the ZEN LITE (ZEISS). The subcellular distribution of CHC (Channel 1, green) was quantified by measuring the line profile of fluorescence intensity from the plasma membrane (marked by Actin staining, channel 2, Magenta) to the nucleus (marked by DAPI staining, channel 3, Cyan). One line was drawn in each cell and data were collected from 30 line profiles (30 cells) for one genotype.

### Figure S6

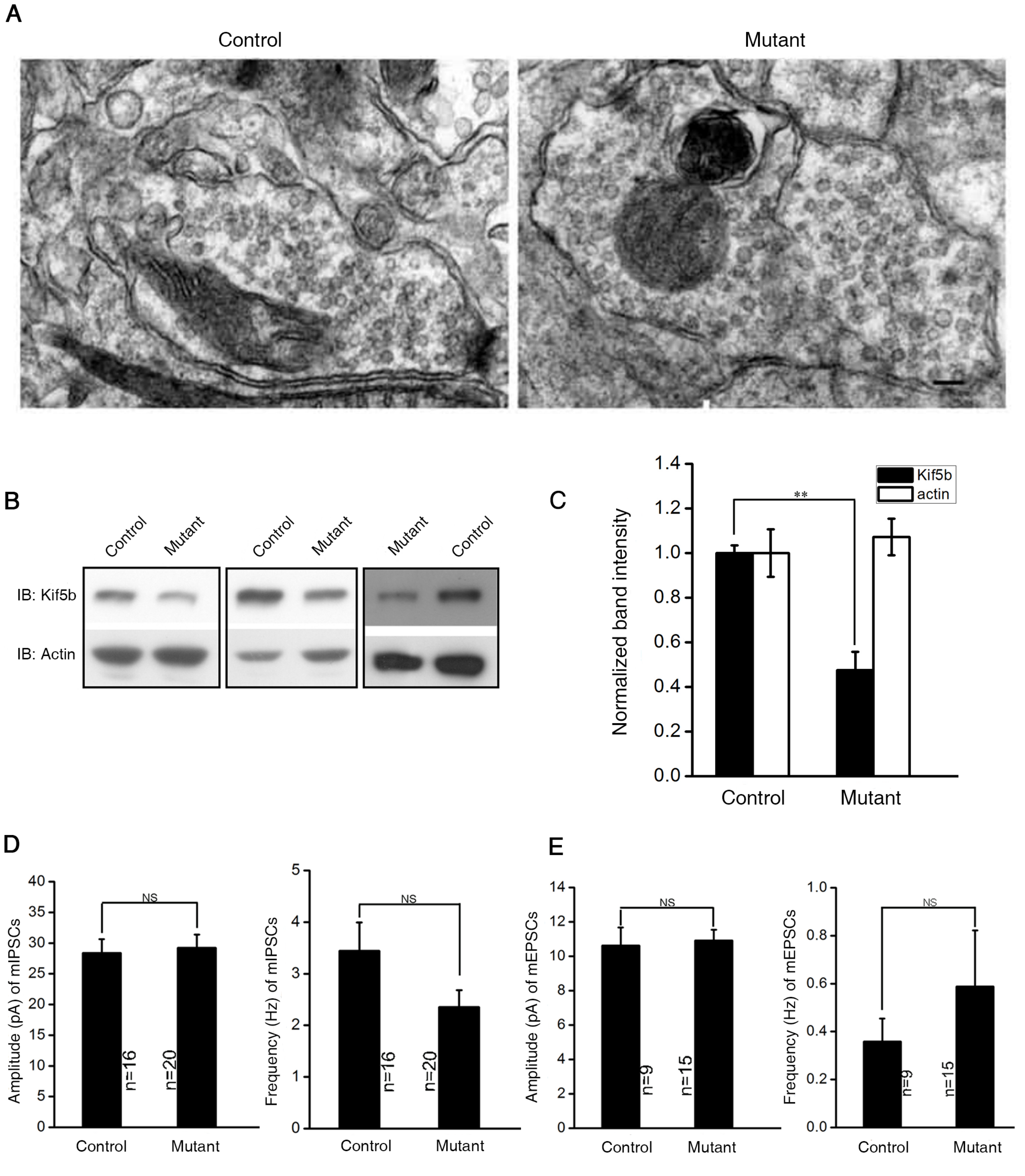

A
Control
Mutant
B
Control
Mutant
Control
Mutant
Mutant
Control
IB: Kif5b
IB: Actin
C
Normalized band intensity
1.4
1.2
1.0
0.8
0.6
0.4
0.2
0.0
Kif5b
actin
**
Control
Mutant
D
Amplitude (pA) of mIPSCs
40
35
30
25
20
15
10
5
0
NS
n=16
n=20
Control
Mutant
Frequency (Hz) of mIPSCs
5
4
3
2
1
0
NS
n=16
n=20
Control
Mutant
E
Amplitude (pA) of mEPSCs
14
12
10
8
6
4
2
0
NS
n=9
n=15
Control
Mutant
Frequency (Hz) of mEPSCs
1.0
0.8
0.6
0.4
0.2
0.0
NS
n=9
n=15
Control
Mutant

**Figure S6. No alteration of synaptic transmission was caused by specific depletion of Kif5b in mouse hippocampus, related to Figure 4.**

(**A**) Electron microscopy examination of synapses revealed no accumulation of coated structures in *kif5b* mutant (right) compared with control (left) mouse brain samples. Scale bar = 100 nm.
(**B**) Representative Western blots of Kif5b in hippocampus of kif5b mutant mice or control littermates. Kif5b expression was remarkably down-regulated by *kif5b* knockout.
(**C**) Quantitative analysis of the indicated bands in left from mutant (n=3) and control (n=3) littermates (right). The levels of Kif5b or Actin in controls were normalized to 1.0, and the levels of the proteins in mutant samples were expressed as the relative to the normalized values. Error bars indicate s.e.m. **P<0.01 as analyzed by Student's t-test.
(**D**) Whole-cell patch-clamp recordings of pyramidal neurons in 20-day-old mouse hippocampal sections were performed. Amplitude (left) and frequency (right) of mIPSCs from cortical sections of 20-day-old control and mutant mice.
(**E**) Amplitude (left) and frequency (right) of mEPSCs from cortical sections of 20-day-old control and mutant mice.

**Table S1. The list of proteins identified in GeL-MS/MS**

| Protein ID | Protein names | Gene names | Log2 LFQ intensity | | | | | | -Log welch p value | MS/MS Count | Mol. weight [kDa] | Sequence coverage [%] |
|---|---|---|---|---|---|---|---|---|---|---|---|---|
| | | | IgG ctrl_1 | IgG ctrl_2 | IgG ctrl_3 | Kif5b _1 | Kif5b _2 | Kif5b _3 | | | | |
| **Q61768** | **Kinesin-1 heavy chain** | **Kif5b** | **15.63** | **13.96** | **16.02** | **27.02** | **25.76** | **26.54** | **2.37** | **208** | **109.55** | **59.6** |
| Q8BGQ1 | Spermatogenesis-defective protein 39 homolog | Vipas39 | 17.54 | 13.47 | 14.59 | 26.17 | 26.08 | 25.80 | 1.90 | 84 | 56.62 | 56.4 |
| Q02819 | Nucleobindin-1 | Nucb1 | 13.29 | 12.96 | 12.80 | 23.99 | 23.80 | 22.99 | 2.99 | 40 | 53.41 | 41.4 |
| Q5UE59 | Kinesin light chain 1 | Klc1 | 14.33 | 14.36 | 14.20 | 25.20 | 24.51 | 24.34 | 3.18 | 96 | 61.63 | 44.3 |
| Q64464 | Cytochrome P450 3A13 | Cyp3a13 | 14.26 | 14.64 | 14.97 | 24.51 | 24.68 | 24.33 | 3.27 | 1 | 57.49 | 3.6 |
| P59016 | Vacuolar protein sorting-associated protein 33B | Vps33b | 15.14 | 14.09 | 14.80 | 24.06 | 23.62 | 23.79 | 2.87 | 47 | 70.53 | 29 |
| D3YXZ3 | Kinesin light chain 2 | Klc2 | 13.40 | 13.95 | 15.18 | 23.11 | 22.56 | 22.33 | 2.34 | 35 | 68.18 | 38.2 |
| Q9DBS5 | Kinesin light chain 4 | Klc4 | 14.61 | 12.83 | 15.78 | 23.00 | 22.34 | 22.20 | 1.92 | 32 | 68.61 | 35.5 |
| Q8BJA3 | Homeobox-containing protein 1 | Hmbox1 | 14.23 | 11.42 | 13.11 | 21.17 | 21.22 | 20.52 | 1.96 | 11 | 47.12 | 20.5 |
| **Q68FD5** | **Clathrin heavy chain 1** | **Cltc** | **20.49** | **10.97** | **20.94** | **25.03** | **24.98** | **24.95** | **0.83** | **131** | **191.55** | **34.6** |
| P17426 | AP-2 complex subunit alpha-1 | Ap2a1 | 13.38 | 12.66 | 15.07 | 20.72 | 20.24 | 20.00 | 1.90 | 14 | 107.66 | 13.4 |
| P84091 | AP-2 complex subunit mu | Ap2m1 | 16.15 | 15.20 | 16.98 | 22.63 | 22.59 | 22.34 | 2.19 | 32 | 49.65 | 31.5 |
| P19253 | 60S ribosomal protein L13a | Rpl13a;Rpl13a-ps1 | 14.66 | 14.30 | 14.08 | 20.97 | 20.71 | 20.37 | 2.84 | 5 | 23.46 | 12.8 |
| P49117 | Nuclear receptor subfamily 2 group C member 2 | Nr2c2 | 14.59 | 13.68 | 14.48 | 20.71 | 20.44 | 20.12 | 2.54 | 18 | 65.24 | 16.8 |
| Q9Z0U1 | Tight junction protein ZO-2 | Tjp2 | 16.21 | 13.61 | 14.61 | 20.73 | 20.13 | 20.31 | 1.72 | 16 | 131.28 | 8.6 |
| Q61548 | Clathrin coat assembly protein AP180 | Snap91 | 13.49 | 13.50 | 14.93 | 19.35 | 19.03 | 19.09 | 2.06 | 7 | 91.85 | 5.3 |
| Q6L8S8 | Beta-1,4-N-acetylgalactosaminyltransferase 3 | B4galnt3 | 13.97 | 12.50 | 15.32 | 19.37 | 18.90 | 18.49 | 1.55 | 2 | 113.51 | 1 |
| P17427 | AP-2 complex subunit alpha-2 | Ap2a2 | 13.73 | 13.57 | 14.79 | 19.58 | 18.96 | 18.27 | 1.93 | 8 | 104.02 | 7.8 |
| O08576 | RUN domain-containing protein 3A | Rundc3a | 14.48 | 12.87 | 16.84 | 19.73 | 19.55 | 18.95 | 1.24 | 7 | 50.02 | 7.6 |
| P28738 | Kinesin heavy chain isoform 5C | Kif5c | 14.27 | 13.37 | 14.24 | 19.27 | 18.08 | 18.39 | 2.01 | 7 | 109.27 | 19.5 |
| D6RIK9 | Protein fantom | Rpgrip1l | 13.64 | 13.41 | 14.99 | 19.27 | 17.86 | 18.25 | 1.69 | 1 | 24.00 | 6.6 |
| Q9Z0E6 | Interferon-induced guanylate-binding protein 2 | Gbp2 | 14.44 | 14.00 | 13.84 | 17.87 | 20.69 | 17.01 | 1.23 | 1 | 66.74 | 1.9 |
| Q9DBG3 | AP-2 complex subunit beta | Ap2b1 | 18.87 | 13.53 | 18.35 | 21.05 | 21.07 | 21.54 | 0.89 | 18 | 104.58 | 11 |
| D3YZG8 | Probable bifunctional methylenetetrahydrofolate dehydrogenase/cyclohydrolase 2 | Mthfd2l | 14.68 | 13.54 | 13.94 | 18.76 | 18.47 | 17.80 | 1.99 | 1 | 36.44 | 3.3 |
| CON__P02538 | | | 19.62 | 14.49 | 15.62 | 19.54 | 20.82 | 21.92 | 0.88 | 35 | 60.04 | 43.6 |
| Q6ZQ06 | Centrosomal protein of 162 kDa | Cep162 | 15.18 | 12.87 | 14.02 | 17.37 | 17.38 | 18.88 | 1.36 | 3 | 160.85 | 0.6 |
| REV__D3Z6B1 | | | 14.86 | 14.59 | 15.30 | 18.28 | 19.19 | 18.28 | 2.00 | 1 | 47.65 | 0 |
| Q3U7K7 | E3 ubiquitin-protein ligase TRIM21 | Trim21 | 14.67 | 12.81 | 14.69 | 18.05 | 17.87 | 17.18 | 1.48 | 6 | 53.33 | 10 |

| | | | | | | | | | | | | |
|---|---|---|---|---|---|---|---|---|---|---|---|---|
| P41105 | 60S ribosomal protein L28 | Rpl28 | 13.11 | 14.62 | 14.41 | 17.08 | 17.86 | 17.73 | 1.66 | 3 | 15.73 | 16.1 |
| Q9CZ44 | NSFL1 cofactor p47 | Nsfl1c | 15.00 | 13.62 | 13.65 | 17.23 | 17.91 | 17.18 | 1.65 | 1 | 40.71 | 4.1 |
| Q3UUI3 | Acyl-coenzyme A thioesterase THEM4 | Them4 | 14.22 | 13.55 | 15.08 | 17.36 | 17.71 | 17.62 | 1.73 | 2 | 26.03 | 16.1 |
| Q8CGY8-2 | UDP-N-acetylglucosamine--peptide N-acetylglucosaminyltransferase 110 kDa subunit | Ogt | 14.48 | 13.99 | 15.08 | 17.95 | 17.52 | 17.03 | 1.73 | 5 | 115.73 | 5 |
| Q9CQF3 | Cleavage and polyadenylation specificity factor subunit 5 | Nudt21 | 14.53 | 15.37 | 15.24 | 18.17 | 18.34 | 17.36 | 1.74 | 6 | 26.24 | 11.9 |
| P27659 | 60S ribosomal protein L3 | Rpl3 | 18.29 | 14.18 | 15.01 | 18.59 | 18.46 | 19.11 | 0.82 | 6 | 46.11 | 6.9 |
| P01630 | Ig kappa chain V-II region 7S34.1 | | 14.24 | 13.93 | 15.46 | 17.48 | 17.47 | 17.25 | 1.58 | 4 | 12.50 | 21.2 |
| P61164 | Alpha-centractin | Actr1a | 18.26 | 14.85 | 14.18 | 18.63 | 18.21 | 18.91 | 0.80 | 8 | 42.61 | 16.8 |
| P01668 | Ig kappa chain V-III region PC 7210 | | 15.29 | 12.24 | 15.60 | 16.83 | 17.65 | 17.01 | 0.90 | 2 | 11.95 | 30.9 |
| Q80YE4 | Serine/threonine-protein kinase LMTK1 | Aatk | 14.03 | 12.94 | 13.70 | 16.91 | 16.13 | 15.69 | 1.52 | 2 | 144.61 | 1 |
| Q9WV60 | Glycogen synthase kinase-3 beta | Gsk3b | 14.10 | 13.34 | 14.50 | 17.40 | 16.05 | 16.50 | 1.44 | 2 | 46.71 | 12.6 |
| Q02257 | Junction plakoglobin | Jup | 14.79 | 13.31 | 14.39 | 17.39 | 16.55 | 16.43 | 1.41 | 10 | 81.80 | 14 |
| Q8QZR5 | Alanine aminotransferase 1 | Gpt | 14.72 | 13.64 | 14.58 | 16.65 | 17.12 | 17.06 | 1.71 | 3 | 55.14 | 4.4 |
| Q9WUX5 | Protein MRVI1 | Mrvi1 | 14.60 | 12.53 | 15.22 | 18.13 | 15.87 | 15.96 | 0.83 | 1 | 97.43 | 1 |
| Q8BFZ9 | Erlin-2;Erlin-1 | Erlin2;Erlin1 | 15.98 | 14.48 | 13.49 | 16.53 | 17.39 | 17.06 | 1.03 | 5 | 37.87 | 16.5 |
| P46471 | 26S protease regulatory subunit 7 | Psmc2 | 13.63 | 12.06 | 14.55 | 15.50 | 15.32 | 16.32 | 1.00 | 2 | 48.65 | 5.3 |
| Q4U2R1 | E3 ubiquitin-protein ligase HERC2 | Herc2 | 13.72 | 14.00 | 15.27 | 16.76 | 17.05 | 15.96 | 1.22 | 2 | 527.45 | 0.8 |
| P14131 | 40S ribosomal protein S16 | Rps16 | 15.34 | 13.62 | 14.65 | 15.73 | 17.12 | 16.94 | 1.05 | 3 | 16.45 | 12.3 |
| Q91XT4 | Protein transport protein Sec16B | Sec16b | 16.05 | 13.80 | 15.12 | 17.04 | 17.13 | 16.68 | 1.01 | 1 | 115.51 | 1.1 |
| Q9EQG3 | Sciellin | Scel | 15.91 | 14.59 | 14.89 | 16.91 | 17.04 | 16.23 | 1.11 | 1 | 72.97 | 1.4 |
| P17897 | Lysozyme C-1 | Lyz1 | 15.07 | 14.39 | 13.77 | 15.54 | 15.87 | 16.20 | 1.13 | 1 | 16.79 | 8.1 |
| P62814 | V-type proton ATPase subunit B, brain isoform | Atp6v1b2;Atp6v1b1 | 16.50 | 16.55 | 17.80 | 13.92 | 16.19 | 14.44 | 0.92 | 4 | 56.55 | 10.6 |
| P80315 | T-complex protein 1 subunit delta | Cct4 | 17.86 | 17.65 | 19.56 | 16.71 | 15.58 | 16.18 | 1.07 | 9 | 58.07 | 12.8 |
| P47738 | Aldehyde dehydrogenase, mitochondrial | Aldh2 | 16.06 | 16.44 | 16.28 | 14.31 | 13.60 | 14.23 | 1.90 | 2 | 56.54 | 4.8 |
| P46660 | Alpha-internexin | Ina | 15.80 | 15.97 | 16.82 | 13.83 | 14.22 | 13.75 | 1.64 | 2 | 55.74 | 5 |
| Q6DFY2 | | Opcml | 16.20 | 16.31 | 16.80 | 13.60 | 14.28 | 14.50 | 1.71 | 2 | 37.16 | 5.9 |
| P35278 | Ras-related protein Rab-5C;Ras-related protein Rab-5A;Ras-related protein Rab-5B | Rab5c;Rab5a;Rab5b | 16.81 | 16.23 | 18.31 | 14.97 | 14.33 | 14.89 | 1.18 | 5 | 23.41 | 22.2 |
| P62492 | Ras-related protein Rab-11A;Ras-related protein Rab-11B | Rab11a;Rab11b | 15.76 | 15.11 | 17.10 | 14.74 | 12.59 | 12.86 | 0.99 | 2 | 24.39 | 11.1 |
| Q3TML0 | Protein disulfide-isomerase A6 | Pdia6 | 16.18 | 16.06 | 17.72 | 14.69 | 14.76 | 12.20 | 0.96 | 5 | 48.69 | 13 |
| P68368 | Tubulin alpha-4A chain | Tuba4a | 16.45 | 15.90 | 17.05 | 12.28 | 14.61 | 13.84 | 1.20 | 3 | 49.92 | 53.6 |
| O35226 | 26S proteasome non-ATPase regulatory subunit 4 | Psmd4 | 16.43 | 16.21 | 16.97 | 14.58 | 13.01 | 13.27 | 1.49 | 1 | 40.70 | 3.7 |
| P18760 | Cofilin-1 | Cfl1 | 17.47 | 17.18 | 18.55 | 14.81 | 13.50 | 14.73 | 1.53 | 6 | 18.56 | 34.3 |

| | | | | | | | | | | | | |
|---|---|---|---|---|---|---|---|---|---|---|---|---|
| F7BX26 | Serine/threonine-protein phosphatase; Serine/threonine-protein phosphatase 5 | Ppp5c | 17.44 | 19.52 | 17.30 | 15.08 | 14.20 | 14.42 | 1.35 | 1 | 54.21 | 2.1 |
| P17751 | Triosephosphate isomerase | Tpi1 | 18.64 | 18.47 | 19.64 | 14.37 | 14.52 | 16.91 | 1.25 | 7 | 32.19 | 23.1 |
| P11983 | T-complex protein 1 subunit alpha | Tcp1 | 16.96 | 16.33 | 19.41 | 13.49 | 13.95 | 14.29 | 1.20 | 7 | 60.45 | 17.6 |
| Q9Z1A1 | | Tfg | 19.95 | 18.22 | 16.85 | 16.43 | 13.83 | 13.72 | 1.00 | 6 | 43.02 | 21.9 |
| Q99PT1 | Rho GDP-dissociation inhibitor 1 | Arhgdia | 17.82 | 17.66 | 18.82 | 14.47 | 13.40 | 14.97 | 1.64 | 4 | 23.41 | 20.6 |
| P97929 | Breast cancer type 2 susceptibility protein homolog | Brca2 | 17.68 | 18.28 | 17.47 | 14.51 | 13.26 | 13.52 | 1.92 | 1 | 370.66 | 0.5 |
| Q8C1B7 | Septin-11 | Sept11 | 18.21 | 18.36 | 18.41 | 13.80 | 14.16 | 14.60 | 2.47 | 5 | 49.69 | 10.7 |
| P52480-2 | Pyruvate kinase PKM | Pkm | 18.37 | 17.07 | 21.18 | 14.64 | 14.39 | 14.55 | 1.16 | 9 | 57.98 | 15.1 |
| P62259 | 14-3-3 protein epsilon | Ywhae | 18.55 | 18.68 | 20.29 | 14.52 | 13.47 | 16.36 | 1.31 | 7 | 29.17 | 25.5 |
| P10639 | Thioredoxin | Txn | 17.11 | 18.21 | 17.70 | 13.70 | 12.52 | 13.62 | 1.90 | 1 | 11.68 | 12.4 |
| P01634 | Ig kappa chain V-V region MOPC 21 | | 17.42 | 18.03 | 18.14 | 14.40 | 12.86 | 13.12 | 1.86 | 2 | 14.90 | 13.2 |
| P18525 | Ig heavy chain V region 5-84;Ig heavy chain V region 914 | | 18.87 | 19.52 | 18.31 | 14.26 | 14.68 | 14.18 | 2.15 | 3 | 12.87 | 23.1 |
| O55131 | Septin-7 | Sept7 | 18.36 | 18.02 | 18.80 | 14.00 | 13.67 | 13.91 | 2.54 | 8 | 50.55 | 19.7 |
| Q9R0T7 | | Try4;Try5 | 21.42 | 21.67 | 20.36 | 19.37 | 14.44 | 14.66 | 1.02 | 3 | 26.27 | 8.1 |
| CON__P02768-1 | | | 18.37 | 18.36 | 18.24 | 12.87 | 13.68 | 12.79 | 2.53 | 2 | 69.37 | 10.3 |
| Q61595 | Kinectin | Ktn1 | 18.51 | 19.09 | 17.79 | 14.00 | 12.65 | 12.92 | 1.96 | 2 | 152.59 | 1 |
| REV__G3UWB5 | | | 19.25 | 19.18 | 18.85 | 13.01 | 13.85 | 14.38 | 2.22 | 1 | 33.27 | 0 |
| P01629 | Ig kappa chain V-II region 2S1.3 | | 19.56 | 19.61 | 19.17 | 13.35 | 14.09 | 14.32 | 2.47 | 1 | 12.22 | 11.6 |
| REV__Q9Z2S7-2 | | | 21.29 | 21.55 | 20.82 | 13.54 | 13.13 | 14.06 | 2.69 | 1 | 9.04 | 0 |

#### Supplemental experimental procedures

##### Reagents, antibodies, and plasmids

Protease inhibitor cocktail (PIC) and protein G-Agrose (Roche, 11719416001), and antibodies against KLC (Chemicon, MAB1616), Kif17 (Sigma, K3638), CHC (BD Biosciences, 610500), Cla (Santa Cruz), Clb (Santa Cruz), Hsc70 (Santa Cruz, sc7298 and Enzo, ADI-SPA-816-D), α-adaptin (BD Biosciences 610502), Dynamin I (Santa Cruz, sc-12724), Syntaxin 6 (BD Transduction Laboratories, 610636), Synaptotagmin (BD Transduction Laboratories, 610433), EEA1 (UPSTATE, 07-292), Auxilin (Santa Cruz, sc104213), Hsp40 (Enzo Life Sciences, ADI-SPA-400-D), beta actin (Sigma, A5316), and VSV-G (Abcam, ab1874) were used. The antibody against Kif5b was previously described (Cui et al., 2011, Wang et al., 2013). Cell penetrating peptides synthesized in GL biochem (Shanghai) Ltd are: TAT-$Kif5b^{891-915}$ peptide GRKKRRQRRRPPQDRKRYQQEVDRIKEAVRSKNMARRG and the control TAT-scramble peptide: GRKKRRQRRRPPQDRKRYQQEVDRIKEAVRSKNMARRG. The GST-fused Kif5b fragments were generated by PCR amplification of pCDNA3.Kif5b plasmid and subcloned into pGEX-4T-1 (Novagen).

##### Cell lines and animals

The cell lines used in this study are Neuro2a neuroblastoma cells and 293T human embryo kidney cells. The $kif5b^{+/-}$ and $kif5b^{floxP/floxP}$ mice were generated by gene targeting as described previously (Cui et al.). To obtain a conditional knockout mouse, $kif5b^{+/-}$ mouse was first crossed with a Camk2a-cre transgenic line(Tang et al., 1999) to generate a $kif5b^{+/-}$; *Camk2a-cre* mouse, which was subsequently bred with $kif5b^{floxP/floxP}$ to get the final conditional knockout mice ($kif5b^{-/floxP}$; *Camk2a-cre*) as well as their littermates ($kif5b^{+/floxP}$). All animal experimentation was approved and performed in accord with the guidelines of the Committee on the Use of Live Animals in Teaching and Research at the University of Hong Kong regarding the care and use of laboratory animals.

##### Immunoprecipitation

Mouse cortices were lysed in lysis buffer (50 mM Tris pH 7.4, 150 mM NaCl, 2 mM EDTA, 1% Triton X-100, PIC), followed by centrifugation at 15,000 × g for 15 min. Supernatants were then incubated with antibodies and protein G-Agarose (Roche). The bound vesicles/proteins were washed several times. Samples were eluted in SDS-PAGE sample buffer at 55ºC, and subjected to SDS-PAGE or Western blot analysis using the indicated antibodies. Enhanced chemiluminescence detection was performed using SuperSignal West Pico reagent (Pierce). Intensities of the bands were quantified by ImageJ (NIH).

##### Liquid chromatography/tandem MS analysis

Biological triplicated samples were applied for electrophoresis using NuPAGE® Novex® 4-12% Bis-Tris Protein Gels. In-gel digestion was performed following an optimized protocol (Shevchenko et al., 2006). Nanoflow electrospray ionization tandem mass spectrometric analysis of peptide samples was carried out using LTQ-Orbitrap Velos (Thermo Scientific, Bremen, Germany) interfaced with Agilent's 1200 Series nanoflow LC system.

##### Pull-down assay

GST-fused Kif5b fragments and Flag-tagged fragments of CHC were expressed in BL21 cells at 18 °C and prepared at 4 °C. The GST-fusion proteins were subsequently immobilized on glutathione-Sepharose beads (GE Healthcare Life Sciences, 17075605) and eluted with 0.2 M glutathione (pH 8.0) for the uncoating reaction or incubated with mouse cortical or bacterial extracts containing flag-tagged CHC fragment in lysis buffer (50 mM Tris pH 7.4, 150 mM NaCl, 2 mM EDTA, 1% Triton X-100, PIC) for the pull-down assay.

##### Immunofluorescence microscopy

Neuro-2a cells cultured on glass coverslips were washed by PBS twice, fixed in 4%

paraformaldehyde in PBS for 10 min, permeabilized in 0.25% Triton X-100 in PBS for 10 min, and incubated in blocking buffer (5% donkey serum in PBST) for 30 min at room temperature. Cells were incubated with indicated primary antibodies overnight at 4 °C, followed by incubation with fluorophore-conjugated secondary antibodies for 1hr at room temperature. Coverslips were then mounted onto glass slides with SlowFade™ Gold Antifade Mountant with DAPI (Life Technologies, S36938). Images were acquired by Zeiss LSM780 confocal laser scanning microscope. Images were processed by ZEN Lite (ZEISS).

**EM analysis of purified CCVs**

For EM analysis, enriched CCVs were fixed with 4% PFA and absorbed into carbon-coated formvar grids. The grids with CCVs for gold staining were stained overnight at 4 °C with control IgG, Kif5b antibody or Kif17 antibody, followed by 1 h incubation of 10-nm gold-conjugated secondary antibody. Negative staining was performed with 2.0% uranyl acetate for 10 sec at room temperature. Electron micrographs were collected from randomly-selected fields using Philips EM 208S (Philips). The images were analyzed by a blind analysis.

**Lentivirus generation and infection**

A modified pLL3.7/U6 promoter vector (a gift from Dr. Dong-Yan Jin) with puromycin resistance was used to express Kif5b shRNA targeting the murine kif5b-encoded mRNA (GenBank accession number for murine Kif5b:NM_008448.3). Two shRNA sequences were selected to target murine Kif5b: 5’- GGACAGATGAAGTATAAATTTCAAGAGAATTTATACTTCATCTGTCC-3’ and 5’-GGCTCTTTCTATTATATCATTCAAGAGATGATATAATAGAAAGAGCC-3’. The control sequence did not target mRNA of any genes: 5'- GACTACCGTTGTATAGGTG TTCAAGAGA CACCTATACAACGGTAGTC-3’. To generate the lentivirus, $2\times10^6$ 293T cells were co-transfected with 10 µg of the specified pLL lentiviral vector and 3.3 µg of each of the packaging vectors (pMD2G-VSVG, pRSV-REV, and pMDL g/p RRE) by the calcium phosphate precipitation method. After 48 h transfection, the supernatant from the transfectant was collected and filtered through 0.45-µm filters (Corning). The medium of the Neuro2a cells was replaced with virus-containing supernatant supplemented with 8 µg/ml polybrene (Sigma, 107689) and incubated for 24 h. Puromycin at a final concentration of 1 µg/ml was added to the culture medium 48 h after transduction.

**Cell-penetrating peptides uptake**

Neuro2a cells were treated with TAT-Kif5b$^{891-915}$ or TAT-scramble peptide diluted to 100 µM in pre-warmed DMEM medium with 10% FBS and incubated at 37 °C for 1 h.

**Statistical analysis**

All data are expressed as mean ± s.e.m. Student's t-test (unpaired) was used to compare two groups ($P$ <0.05 being considered significant).
